# Ultraviolet Photodissociation of tryptic peptide backbones at 213 nm

**DOI:** 10.1101/2020.03.25.008326

**Authors:** Lars Kolbowski, Adam Belsom, Juri Rappsilber

## Abstract

We analyzed the backbone fragmentation behavior of tryptic peptides of a four protein mixture and of *E. coli* lysate subjected to Ultraviolet Photodissociation (UVPD) at 213 nm on a commercially available UVPD-equipped tribrid mass spectrometer. We obtained 15,178 high-confidence peptide-spectrum matches by additionally recording a reference beam-type collision-induced dissociation (HCD) spectrum of each precursor. Type a, b and y ions were most prominent in UVPD spectra and median sequence coverage ranged from 5.8% (at 5 ms laser excitation time) to 45.0% (at 100 ms). Overall sequence fragment intensity remained relatively low (median: 0.4% (5 ms) to 16.8% (100 ms) of total intensity) and remaining precursor intensity high. Sequence coverage and sequence fragment intensity ratio correlated with precursor charge density, suggesting that UVPD at 213 nm may suffer from newly formed fragments sticking together due to non-covalent interactions. UVPD fragmentation efficiency therefore might benefit from supplemental activation, as was shown for ETD. Aromatic amino acids, most prominently tryptophan, facilitated UVPD. This points at aromatic tags as possible enhancers of UVPD. Data are available via ProteomeXchange with identifier PXD018176 and on spectrumviewer.org/db/UVPD_213nm_trypPep.

## INTRODUCTION

The spread of Ultraviolet Photodissociation (UVPD) as an alternative fragmentation method in mass spectrometry has increased substantially in recent years [1]. UVPD uses photons to achieve bond dissociation, which is a fundamentally different mechanism from the widely used collisional-activation based (collision induced dissociation (CID) [2] and beam-type CID (HCD) [3]) and electron-based (electron transfer dissociation (ETD) [4] and electron capture dissociation (ECD) [5]) methods. Photons are typically produced by lasers at specific wavelengths over varying excitation times tapping into a plethora of different use cases and possible applications [6]. Coupling lasers to mass spectrometers often requires highly specialized, custom instrumentation, with important safety considerations that must be met. A recent key update here has been the release of a commercially available 213 nm solid-state laser coupled to a tribrid mass spectrometer. This has opened up the technology to be used by a wider user base outside of specialized labs who have implemented UVPD by making custom modifications to their instruments.

So far 213 nm UVPD has mainly been used in topdown proteomics. For this, Fornelli et al. [7] have recently conducted a thorough performance evaluation. Studies on peptide fragmentation with UVPD at 213 nm have been conducted on in-house modified instruments, only looking at a very low number of peptides and very specific bond cleavages. Girod et al. described specific fragmentation behavior of proline containing peptides in their study on 4 synthetic peptides subjected to a long (1000 ms) 213 nm UVPD excitation time [8]. Mistarz et al. investigated the fragmentation behavior of a single triply protonated 12-mer peptide at 600 ms laser excitation time for localizing sites of backbone deuteration [9]. More recently Talbert and Julian have carried out a study on initiating bond-selective fragmentation using 213 nm UVPD on 13 peptides [10].

In this study we investigated the 213 nm UVPD fragmentation behavior of a large number of tryptic peptides in bottom-up proteomics experiments, using a well-defined four protein mix as well as an E. coli lysate. We deployed a systematic Dual MS2 acquisition scheme, utilizing reference HCD spectra for identification and to obtain UVPD peptide-spectra matches (PSMs) independent of UVPD fragmentation efficiency. We characterized peptide backbone ion-types present and compared the sequence coverage and the sequence fragment intensity ratio across a range of UVPD excitation times and other fragmentation techniques. As the speed of acquisition plays a crucial role on the number of possible identifications [11] we investigated excitation times comparable to typical ETD reaction timescales [12] to acquire more practical data points. Finally, we probed our data to find precursor dependent properties that facilitate good UVPD fragmentation.

## EXPERIMENTAL METHODS

### Materials/Reagents

Human serum albumin (HSA), ovotransferrin (chicken) and myoglobin (equine) were purchased from Sigma-Aldrich (St. Louis, MO). Creatine kinase (rabbit) was purchased from Roche (Basel, Switzerland).

### Sample preparation

HSA, ovotransferrin, myoglobin and creatine kinase were dissolved in 8 M urea with 50 mM ammonium bicarbonate to a concentration of 2 mg/mL each. Proteins were reduced by adding dithiothreitol at 2.5 mM followed by incubation for 30 min at 20 °C. Samples were derivatized using iodoacetamide at 5 mM concentration for 20 min in the dark at 20 °C, diluted 1:5 with 50 mM ammonium bicarbonate and digested with trypsin (Pierce Biotechnology, Waltham, MA) at a protease-to-protein ratio of 1:100 (w/w) during a 16-h incubation period at 37 °C. Digestion was stopped by adding 10% TFA at a concentration of 0.5%.

E. coli K12 cells were cultured in LB medium, harvested at OD 0.6, pelleted, frozen using liquid nitrogen and stored at −80 °C. Cell pellets were then resuspended in ice-cold lysis buffer (20 mM HEPES, 150 mM NaCl, pH 7.5 and protease inhibitors (Roche)) and sonicated on ice at 30% amplitude, 30 s on/off for 10 cycles (total time 5 mins), using a Branson Digital Sonifier. Liquid was collected in a centrifuge tube (upper foaming with DNA proteins was discarded) and sample was clarified by centrifugation at 15,500 rpm for 30 mins. Protein concentration was measured using the Pierce BCA protein assay. The soluble lysate was then run into the first centimeter of an SDS-PAGE gel, the protein band excised and subjected to an in-gel digestion protocol: reduction in 20 mM DTT for 30 min at room temperature, alkylation with 55 mM iodoacetamide for 30 min at room temperature in the dark, and digestion with 12.5 ng/μL trypsin overnight at 37 °C.

Digests for all proteins were cleaned up using the StageTip protocol [13]. Peptides were eluted using 80% v/v ACN, 0.1% v/v TFA, partially evaporated using a Vacufuge Concentrator (Eppendorf, Germany) to <5% ACN, and resuspended in mobile phase A (0.1% formic acid) prior to mass spectrometry analysis.

### Data acquisition

Samples were analyzed using an UltiMate 3000 Nano LC system coupled to an Orbitrap Fusion Lumos Tribrid mass spectrometer equipped with an EasySpray source and UVPD module (Thermo Fisher Scientific, San Jose, CA) comprising a solid-state Nd:YAG laser head (Cry-LaS GmbH) that generates a pulsed 213 nm beam (fifth harmonic). Mobile phase A consisted of 0.1% formic acid in water, mobile phase B of 80% acetonitrile, 0.1% formic acid, and 19.9% water. Peptides (1 μg eq.) were loaded onto a 500 mm C-18 EasySpray column (75 μm ID, 2 μm particles, 100 Å pore size) and eluted using the following non-linear gradient at 300 nl flow rate: 2-4% (1 min); 4-6% B (2 min); 6-37.5% B (97 min); 37.5-42.5% B (10 min); 42.5-47.5% B (5 min). MS1 spectra were acquired in the Orbitrap using a scan range from 300-1500 *m/z* at 120,000 resolution. AGC Target was set to 8 × 105 and maximum injection time to 50 ms. Filters used were: Peptide monoisotopic precursor selection (MIPS), intensity threshold of 5 × 104, precursor charge states 2-7+ and dynamic exclusion of 60s. MS2 spectra were acquired using top speed with a 3 s cycle time at 30,000 Orbitrap resolution. Precursors were isolated using the Quadrupole with an isolation window of 1.4 *m/z*. MS2 AGC target was set to ‘Standard’ using the ‘Auto’ maximum injection time mode. For each precursor two MS2 scans were acquired (Dual MS2), one with HCD at NCE 30%, the other utilizing UVPD at 213 nm with varying laser excitation times between experiments (5, 10, 15, 20, 30, 40, 50 and 100 ms). As reference Dual MS2 with HCD-HCD, HCD-ETD and HCD-EThcD (Supplemental activation with 30% NCE) were acquired. For the ETD experiments calibrated charge-dependent reaction times were used. For each Dual MS2 method two replicas were acquired, resulting in a total of 22 acquisitions per sample.

### Data Analysis

Raw files were preprocessed using a custom python script (https://github.com/Rappsilber-Laboratory/preprocessing). Preprocessing included conversion to MGF format, *m/z* recalibration of both precursor and fragment peaks as well as splitting the primary (HCD) and secondary (UVPD, ETD, EThcD or HCD) MS2 spectra into separate MGF files. The primary MS2 spectra were denoised with the default MS2Denoise settings in MSconvert [14] and then analyzed with xiSEARCH [15] using the following settings: MS accuracy, 3 ppm; MS2 accuracy, 15 ppm; missing monoisotopic precursor peaks, 2; enzyme, trypsin; maximum missed cleavages, 4; maximum number of modifications, 3; ions: precursor, b- and y-type; modifications: carbamidomethylation on Cysteine as fixed and oxidation on Methionine as variable modification; False Discovery Rate was estimated separately for each fragmentation parameter set (2 acquisition replica per fragmentation parameter) using xiFDR [16] on unique PSM level to 1% for the 4 PM and 0.1% for the E. coli lysate. Additionally, we excluded PSMs that were not identified in at least 6 of the 11 different fragmentation parameter set duplicates. This minimizes the number of false positives in our data set because of the improbability of any given random match to occur in more than half of the experiments. From the secondary peak list files the corresponding scans to the primary HCD PSMs were extracted and subsequently annotated with pyXiAnnotator (https://github.com/Rappsilber-Laboratory/pyXiAnnotator/) using the identifications from their primary counterparts. Secondary spectra were annotated with peptide, a-, b-, c-, x-, y- and z-type ions with a maximum tolerance of 15 ppm. Additionally, a hydro-gen-loss was defined and the missing monoisotopic fragment peak feature was used to enable matching of ions with absence or presence of an extra hydrogen respectively (e.g. y—, a+).

The mass spectrometry raw files, peak lists, search engine results and FASTA files have been deposited to the ProteomeXchange Consortium via the PRIDE [17] partner repository with the dataset identifier PXD018176 and 10.6019/PXD018176. Additionally, we have made all annotated spectra available through xiSPEC [18] on spectrumviewer.org/db/UVPD_213nm_trypPep.

## RESULTS/DISCUSSION

### Dual MS2 data acquisition

We aimed to evaluate UVPD efficiency of peptides in tryptic digests at different excitation times. For some settings and peptides the resulting MS2 spectra themselves may not allow unambiguous identification of the peptide due to poor fragmentation efficiency. We therefore designed a Dual MS2 acquisition approach, i.e. acquiring two MS2 spectra for each precursor: one fragmented with HCD and then another fragmented with UVPD (alternatively ETD, EThcD or HCD as controls). This allowed us to use the well-established collisional fragmentation method to reliably identify precursors, and then annotate and evaluate the UVPD spectra with those identifications, even when the extent of peptide fragmentation in a UVPD spectrum alone did not enable unambiguous fragment-based identification (Figure 1).

**Figure 1.**
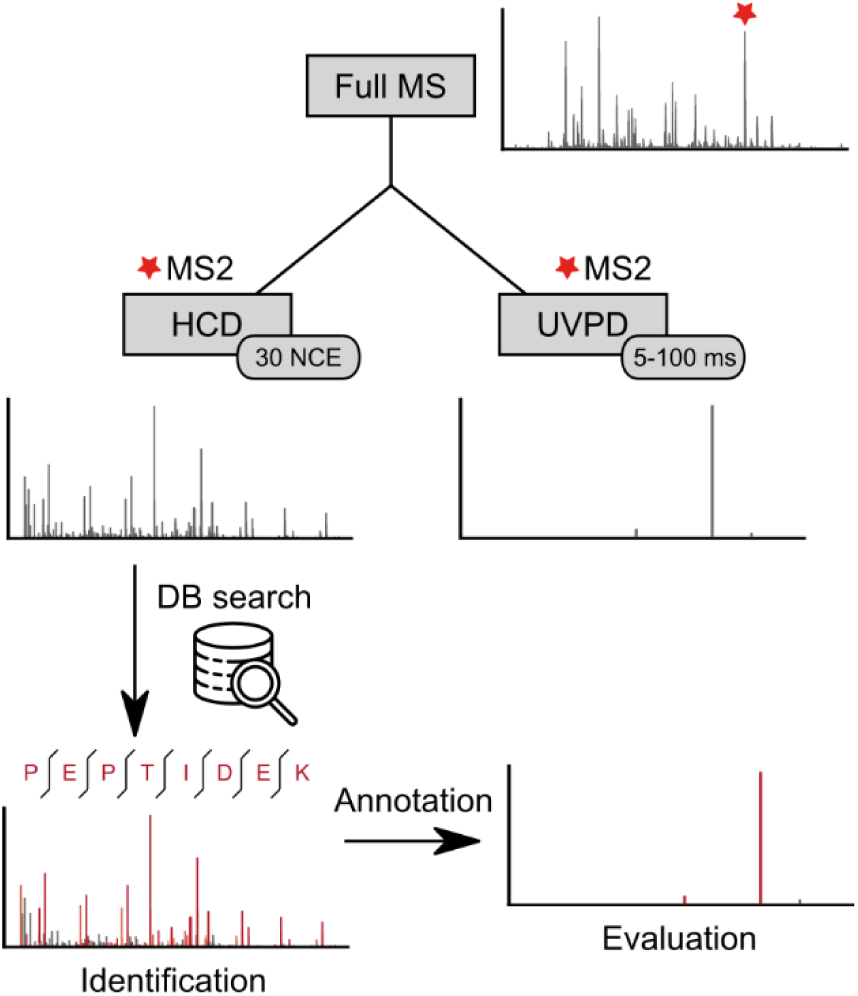
Dual MS2 workflow. For each selected precursor two MS2 spectra are acquired, one with HCD and one with UVPD fragmentation. The HCD spectrum is used for identification via database search. The identified peptide is then used to annotate the UVPD spectra, allowing evaluation even when the extent of peptide fragmentation in a UVPD spectrum alone does not enable unambiguous fragment-based identification.

Using the HCD scans for identification gave us a high number of PSMs over all tested conditions (Figure S1). This becomes especially apparent for the short excitation time acquisitions of the more complex E. coli sample. From the 5 ms acquisitions 17,536 UVPD spectra could be evaluated using identification from the corresponding primary HCD spectra, whereas relying on UVPD alone would have only yielded 1,047 PSMs. Furthermore, analyzing the subset of UVPD spectra that led to peptide identifications would have introduced a strong bias towards well fragmenting peptides. The HCD-HCD control samples led to almost identical numbers of PSMs from both the reference HCD as well as from the secondary HCD control scan, showing that the quality of the second spectra is not adversely affected and thus demonstrating the viability of this approach.

To further improve the quality of our data we applied the additional restriction to include only PSMs that were seen in more than half of our acquired fragmentation parameter sets. This rather conservative selection of PSMs (MS1 plus MS2 HCD fragment-based identification with stringent FDR cutoff and the additional requirement mentioned above) combines the benefits of using a real-world sample from a protein digest while approaching the confidence level of a synthetic peptide library.

### Fragmentation analysis of different UVPD excitation times

We investigated the UVPD-induced peptide fragmentation behavior of tryptic digests from a lower complexity sample consisting of four model proteins (4 PM) and a complex E. coli lysate sample and compared it with HCD, ETD as well as EThcD fragmentation.

Our dataset consists of 15,178 unique precursors (unique sequence and charge state combination; 4 PM: 667, E. coli: 14,511; Table S1). These originate from 11,663 unique peptides (4 PM: 384, E. coli: 11,279) subjected to UVPD at 213 nm with varying laser excitation times (5-100 ms). We analyzed both datasets separately to test if we can reproduce the findings from our more complex E. coli lysate sample also in the small well-defined 4 PM sample. The evaluation of fragmentation characteristics from both datasets led indeed to very similar results (compare Figure 2 & Figure S2). Our analysis found that UVPD at 213 nm results in all possible types of backbone fragmentation ions in tryptic peptides, with a-, b- and y-type ions being the most prominent over all excitation times (Figure 2A). Ion counts were normalized per peptide by length for ease of comparison. Ion counts increase with increasing excitation time and start to level off at the maximum acquired time for all ion types. Doubling the excitation time from 50 to 100 ms resulted only in a very small increase in fragment count.

**Figure 2.**
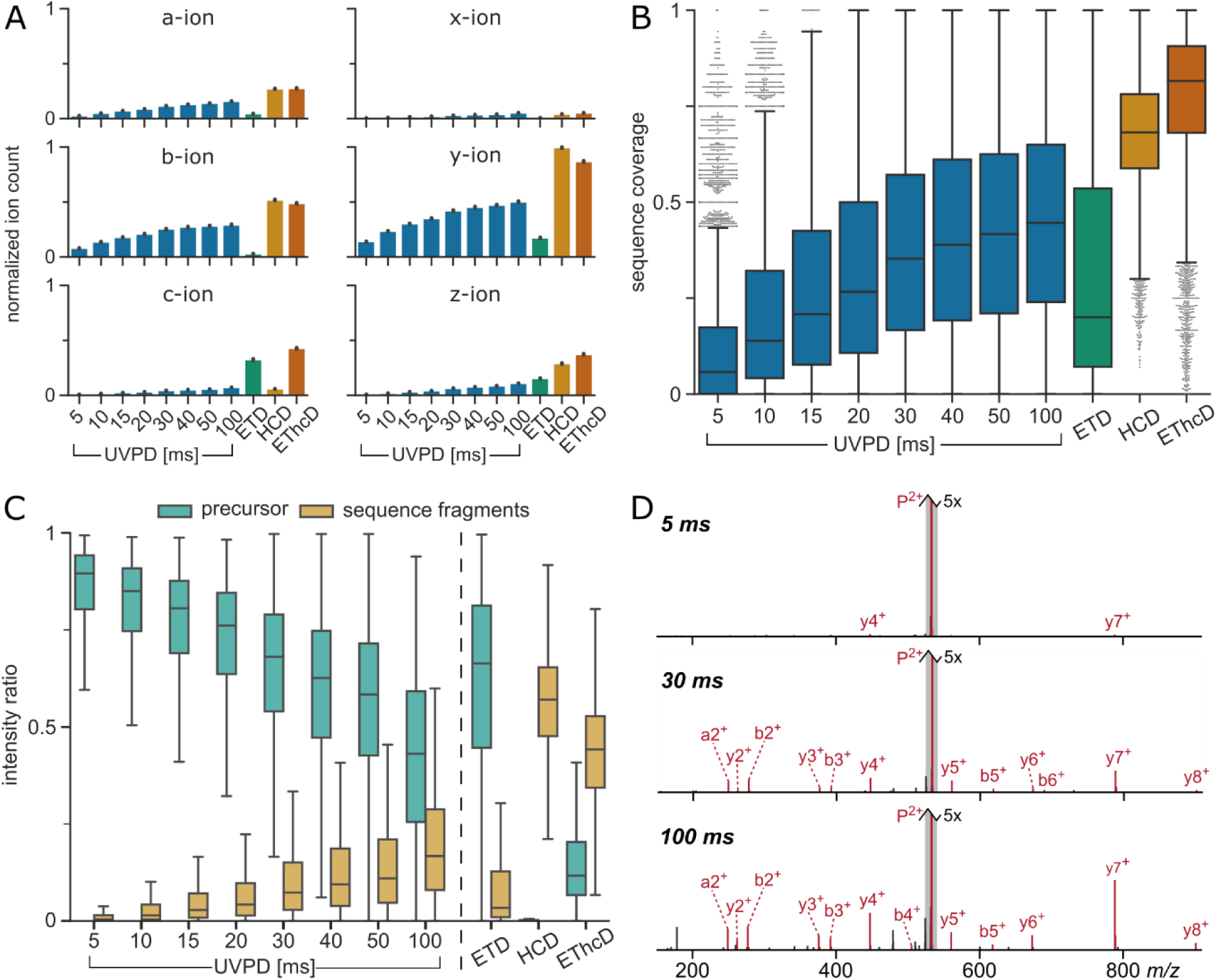
Fragmentation analysis of peptides subjected to different UVPD excitation times and ETD, HCD and EThcD for comparison. Data from *E. coli* dataset. **(A)** Bar plots showing the sequence fragment ion type counts (normalized by peptide length). Error bars reprersent 0.95 confidence intervals. **(B)** Box plots showing sequence coverage of precursors. **(C)** Box plots of MS2 intensity ratios of remaining precursor and sequence fragments to total MS2 intensity. In all box plots whiskers extend to 1.5 interquartile range past the low and high quartiles. **(D)** Example MS2 spectra of YLDLIANDK (charge 2+) subjected to 5, 30 and 100 ms of UVPD. All three spectra are zoomed on the y-axis (5x). Annotated fragment peaks are shown in red, the unfragmented precursor is highlighted with grey outline.

We calculated the sequence coverage for our PSMs conservatively, as the ratio of matched N-terminal and C-terminal sequence fragments to the number of theoretically possible sequence fragments, i.e. 100% sequence coverage would mean detection of at least one fragment from the N-terminal (a, b or c) and one from the C-terminal series (x, y or z) between all amino acid resirdues of a peptide. Sequence coverage of UVPD was low at 5 ms (median: 5.8%) but increased steadily with excirtation time to a median of 45.0% for 100 ms. While still on the low side compared to the HCD at 68.2% and EThcD 81.6%, UVPD started to give better median serquence coverage than ETD with excitation times higher than 15 ms. The low median sequence coverage from ETD is partly due to its dependency on higher charge states (median sequence coverage for charge 2+: 12.5% vs. 41.2% for ≥3+).

Notably even for the lowest UVPD excitation time, there are some precursors that fragmented reasonably well (showing a sequence coverage > 50%) while others showed almost no fragmentation even at 100 ms. This suggested that precursor dependent properties play an important role in UVPD. Also, comparing the last two UVPD data points showed that doubling the excitation time had only a marginal effect on the resulting serquence coverage.

As a way of assessing the fragmentation efficiency we calculated the ratio of the sum of all sequence fragment isotope cluster peak intensities to the total MS2 intensity. This sequence fragment intensity ratio remained low over all tested UVPD excitation times, ranging from a median of 0.4% at 5 ms to 16.8% at 100 ms (Figure 2C).

In contrast to the ion count and the sequence coverage, doubling the excitation time from 50 to 100 ms had a visible effect on the fragmentation efficiency. In conclursion, extending excitation times beyond a precursor-dependent threshold led to higher fragment intensity but few additional fragments.

The high remaining precursor intensity revealed that the low sequence fragment intensity was due to low fragmentation efficiency, similar to what has been described for ETD [19]. This can be seen in the representartive spectra where even at 100 ms UVPD the unfragrmented precursor is the most intense peak by far (Figure 2D). In the HCD spectra on the other hand a median of 57.0% of MS2 intensity stems from sequence fragments with almost no leftover precursor. This suggests that here the remaining intensity most likely stems from interrnal fragmentation events, commonly seen in HCD [20].

Next, we checked the complementarity between the HCD and the secondary scans by adding the annotated fragments from the secondary MS2 scan to its primary scan and recalculating the sequence coverage. In Figure 3 the gain in sequence coverage for each method is plotrted. Note that the combination of two consecutive HCD scans on the same precursor already resulted in an avrerage ∼4% gain and should therefore be considered as a baseline. UVPD with excitation times > 20 ms showed sequence coverage gains above this baseline, but the gain seemed to level off at 40 ms around 7%. Both ETD as well as EThcD were favorable over UVPD in terms of sequence coverage complementarity to HCD.

**Figure 3.**
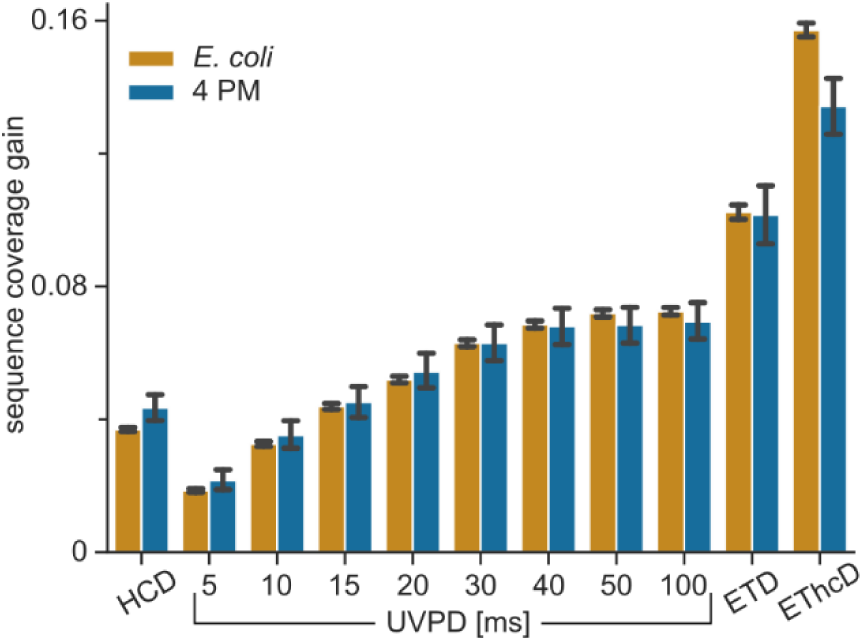
Sequence fragment complementarity of acquired fragmentation methods to the HCD reference spectra. Plotted is the sequence coverage gain for all matches from the *E. coli* and 4 PM datasets when combining annotated fragments from the secondary MS2 scan (HCD, UVPD 5-100 ms, ETD or EThcD) with the refrerence HCD scan versus the reference HCD scan alone.

### Influence of charge state on UVPD

In search of precursor properties that influenced UVPD fragmentation, we first looked at the precursor charge state. We split our PSMs by charge state and evaluated the correlation to the sequence coverage by linear regression fitting (Figure 4A). In contrast to ETD, a higher charge state showed negative correlation with sequence coverage over all tested laser excitation times. The charge density (precursor charge divided by the peptide amino acid length) on the other hand seemed to be an important influence on UVPD fragmentation. A high charge density correlated with high sequence covrerage independent of excitation time (Figure 4B). The influence of the charge density became even more aprparent when comparing spectra of the same peptide that were acquired in multiple charge states. Figure 4 panels C-F show spectra of the peptide QEPER-NECcmFLQHKDDNPNLPR fragmented in charge states 2+, 3+, 4+ and 5+. In the spectrum with the lowest charge state only 4 sequence fragments are visible, i.e. 10% sequence coverage. The sequence coverage inrcreased to 17.5% for charge state 3+ (7 sequence fragrments) and charge state 4+ already showed 27 serquence fragments, resulting in 42.5% sequence coverage. Finally, the highest detected charge state for this peptide (5+) presented a rich fragmentation spectrum with high sequence coverage (67.5%), albeit fragmentartion efficiency still remains relatively low.

**Figure 4.**
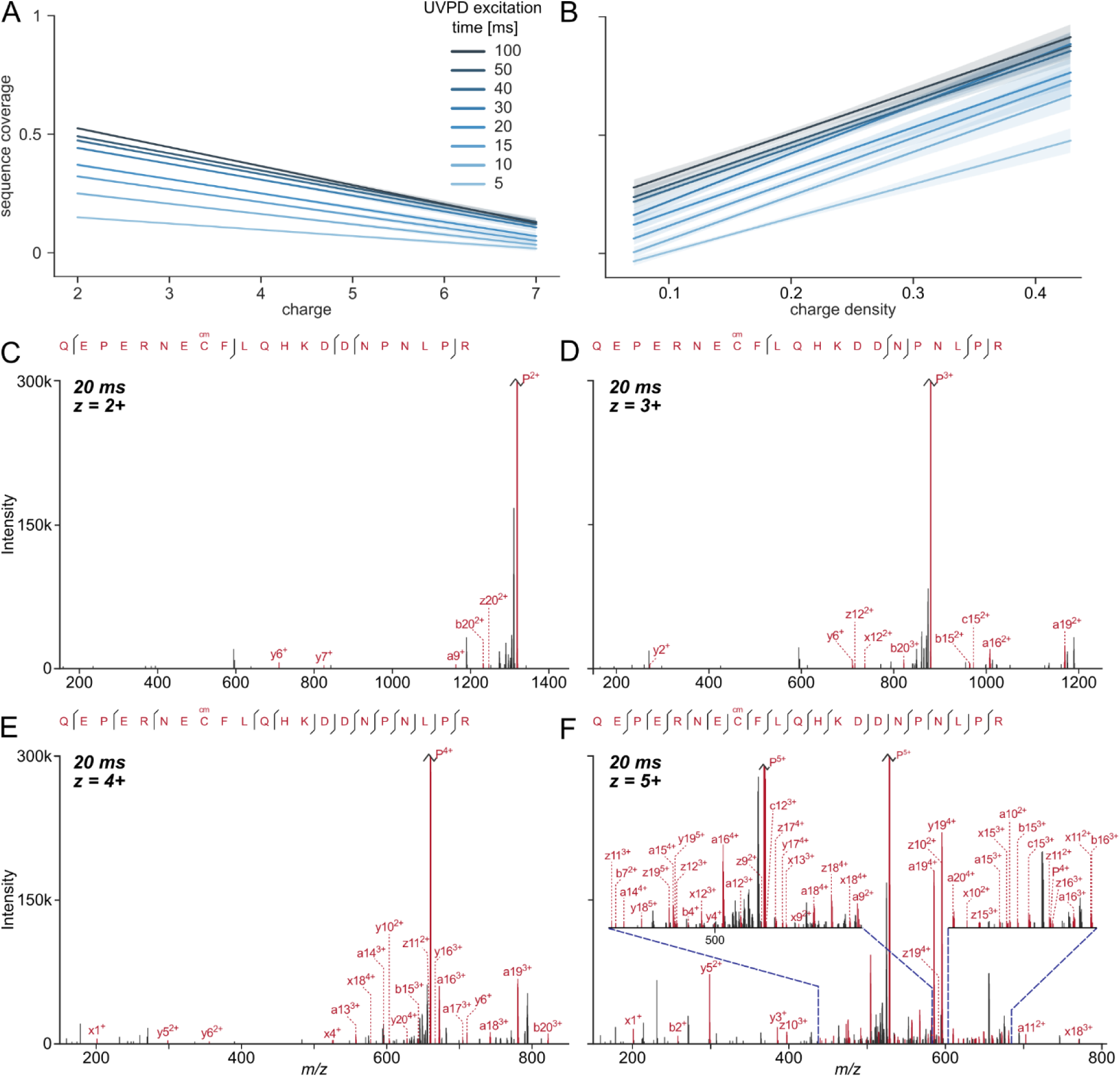
Influence of charge **(A)** and charge density **(B)** on the sequence coverage in UVPD experiments with varying laser excitation times. Linear regression model fit of all analyzed PSM data with 0.95 confidence interval shown as translucent bands around lines. **(C-F)** Example fragmentation spectra of peptide QEPERNECcmFLQHKDDNPNLPR (Ccm stands for carbamidomethylated Cysteine) subjected to 20 ms UVPD in charge states 2+, 3+, 4+ and 5+. Spectra are zoomed in on the y-axis to 300k intensity for visibility and the remaining unfragmented precursor peaks are cut off.

A possible interpretation of the charge density derpendence of UVPD could be incomplete dissociation through noncovalent interactions between newly-formed fragments, a phenomenon which has been previously described for electron-based fragmentation methods [21, 22]. In the case of ETD, several supplemental activation techniques have been developed to address this drawrback using e.g. collision-based activation (ETciD/EtcaD and EThcD [19]) or infrared excitation [24]. Recently, Halim et al. have reported increased sequence coverage in a top-down proteomics experiment through simultanerous and consecutive irradiation using UV (213 nm) and IR (10.6 μm) lasers, coined high-low photodissociation (HiLoPD) [25]. Unfortunately however, thus far there is no commercial instrument available that contains both a UV and an IR laser source. Furthermore, the combinartion of UVPD and collision-based activation is not possirble with the current version of the instrument control software.

### Influence of amino acid composition on UVPD

The availability of chromophores is crucial for absorbring the energy provided by photons without which UVPD will not take place. While for shorter wavelengths like 157 nm almost all bonds can act as chromophores, 213 nm is already at the absorption threshold for smaller chromophores like amide bonds [10]. Talbert and Julian found that 213 nm can give rise to both bond specific as well as nonspecific dissociation depending on laser power and excitation time as well as molecular composirtion. The molecular composition and structure of peprtides is defined by their amino acid composition. We analyzed the cleavage propensity on UVPD for all backbone ion types. The heatmaps in Figure 5 show the relartive likelihood for the detection of a backbone cleavage product between two amino acids. Propensities were calculated by dividing the observed occurrences of a fragmentation site by the theoretical possible values for all combinations of amino acids N- and C-terminal of the site. For ease of comparison, ratios were normalized by the mean for each product ion type and then logarithrmized (base 2). Amino acid pairs between which cleavrage occurs preferentially, differed between the different ion types, but overall the aromatic amino acids and prorline showed hotspots for increased cleavage propensity. Similar to what has been described by Oh et al. [26] for 266 nm and by Fornelli et al. [7] for 213 nm top-down proteomics, we found an increased frequency of fragrmentation events adjacent to aromatic amino acids. Type a- and x-ions and, to lesser extent, also b-, y- and z-ions showed preference for cleavage N-terminal of phenylalanine. Interestingly, c- and z-type cleavage ocrcurred with largely increased frequency N-terminal of tryptophan. The most prominent ion types (a, b and y) all occur with increased frequency with a proline, C-terminal of the fragmentation site. This is in line with findings from Girod et al. [8], who have shown C-C and C-N bond actirvation close to proline residues following 213 nm excitartion of the proline containing peptides and was also seen by Fornelli et al. [7] in top-down analysis.

**Figure 5.**
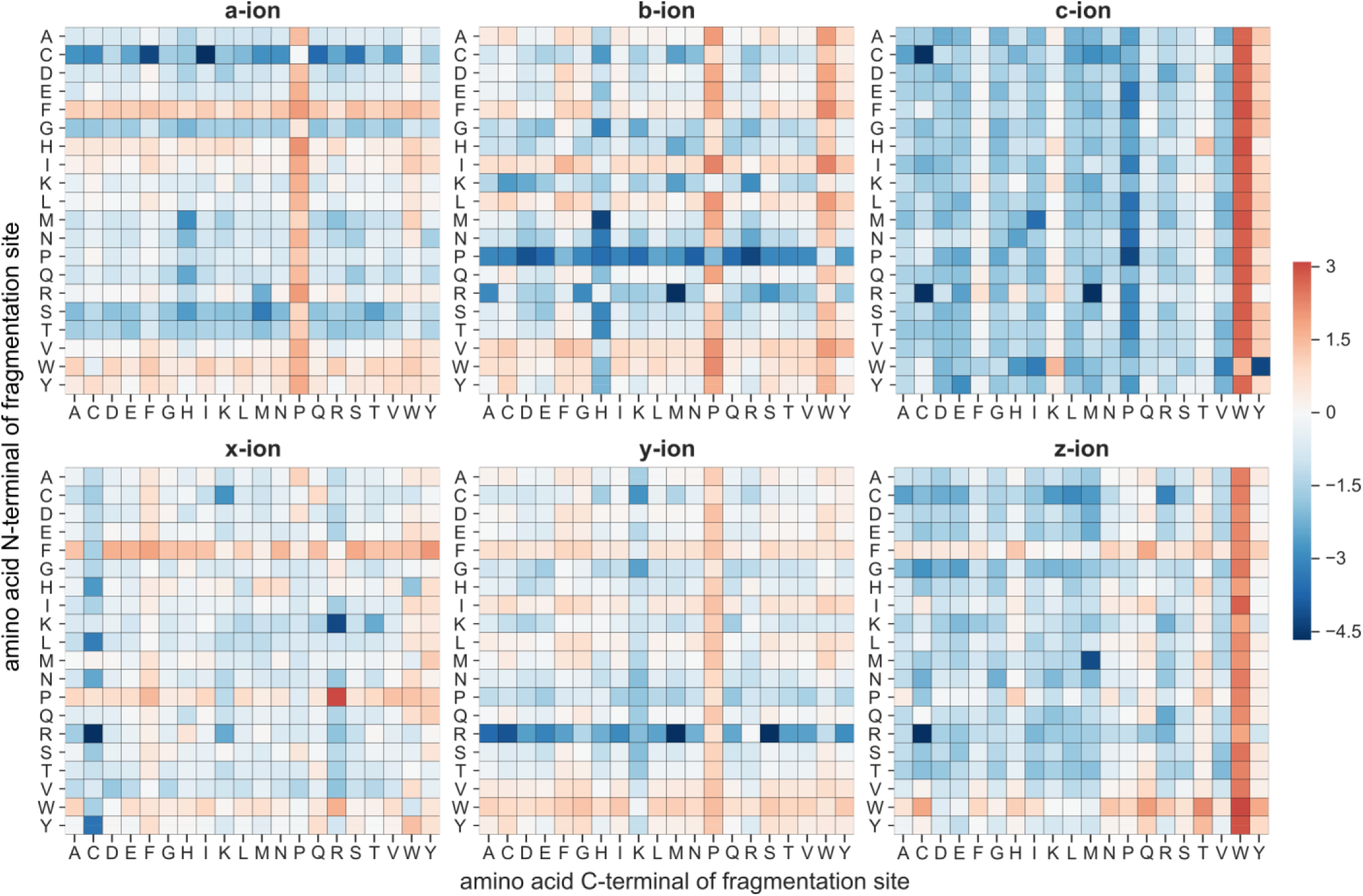
Cleavage propensities of backbone fragment ion types for UVPD. Heatmaps show the ratio of detected versus theoretical cleavrage events for all amino acid combinations that can occur N- and C-terminal of the fragmentation site. Values were normalized by the mean for each ion type and then logarithmized to base 2. White represents average cleavage propensity for this ion type, blue less and red higher than average occurrence.

To systematically check the influence of the amino acrid composition on fragmentation of the whole peptide, we compared the mean amino acid composition of those peptides that fragmented well under UVPD, with those that showed poor fragmentation over all acquired excitartion times. We looked at the sequence coverage as well as the sequence fragment intensity ratio and divided our data into terciles for both those metrics. We calculated the average frequency for each amino acid in the upper and lower tercile and compared the fold-change. This showed which amino acids were disproportionately prevalent in UVPD-susceptible peptides (Figure 6A). Peptides with high sequence coverage had a significantrly increased frequency of tryptophan and phenylalanine. Looking at the sequence fragments intensity ratio, peprtides in the upper tercile exhibited increased ratios of aromatic amino acids (most prominently again tryptorphan). Interestingly, while a higher proline ratio seemed to translate into intense sequence fragments it did not promote high sequence coverage, suggesting that proline only leads to efficient cleavage of specific bonds (adjacent, see Figure 5). Figure 6 panels B and C show example spectra of two peptides at the same charge state and with similar *m/z* both subjected to 20 and 100 ms UVPD. The peptide in panel B contains both a tryprtophan and a phenylalanine and already shows almost complete sequence coverage at 20 ms, but still an inrtense remaining precursor peak. At 100 ms the precurrsor is almost completely fragmented. The peptide in panel C on the other hand contains no aromatic amino acids or proline. There is a complete absence of fragrment peaks at 20 ms and only very little low intense fragment peaks are present in the 100 ms spectrum.

**Figure 6.**
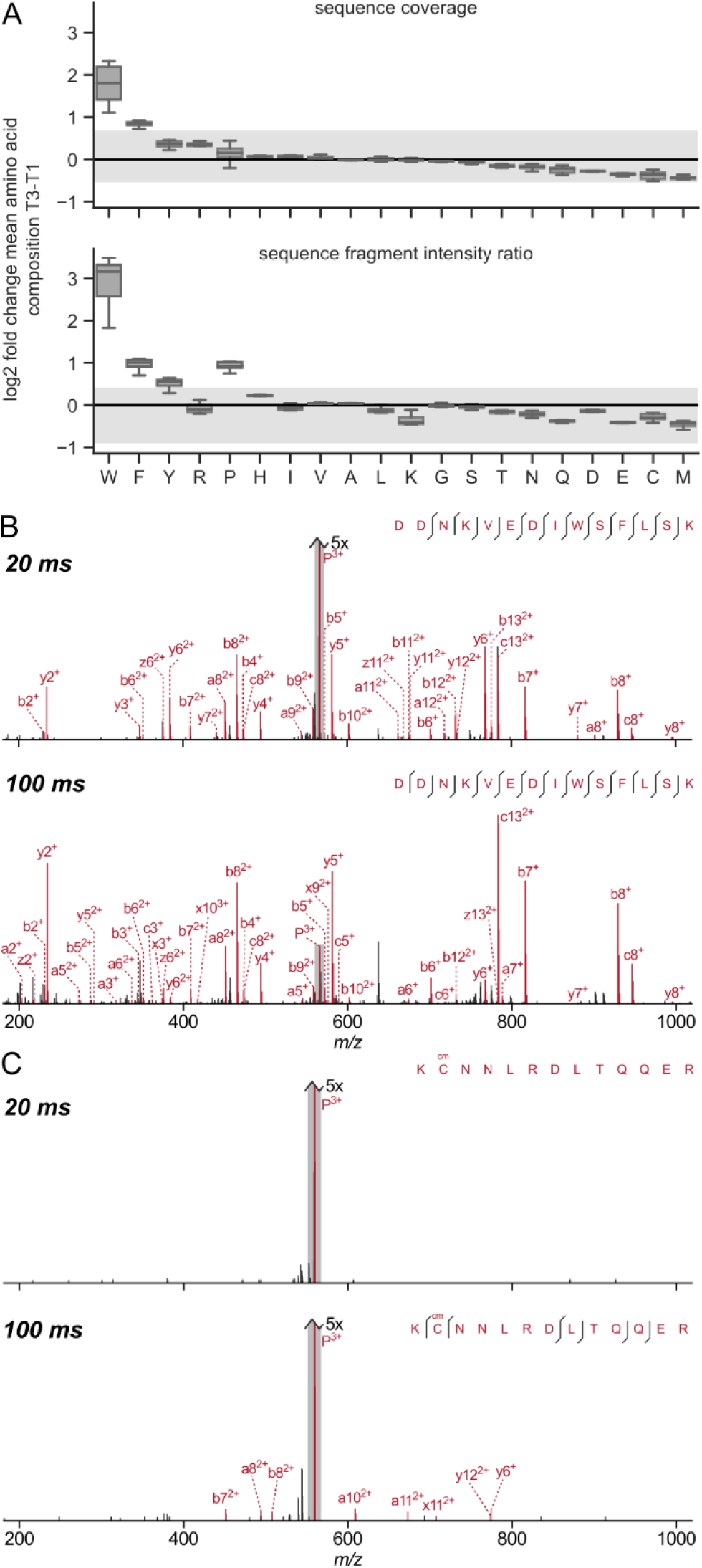
Influence of amino acid composition on peptide back-bone fragmentation for UVPD. **(A)** The boxplots show the log2 fold change of the mean amino acid composition of PSMs in the upper tercile (T1) against the lower tercile (T3) in terms of serquence coverage and sequence fragment intensity ratio, respecrtively. Data from the *E. coli* dataset was considered for the boxrplots. The spread of the respective HCD data is shown in grey as a reference. **(B-C)** Example spectra of two peptides with similar *m/z* and same charge state both subjected to 20 and 100 ms UVPD. DDNKVEDIWSFLSK containing both tryptophan and phenylalarnine **(B)** and KCcmNNLRDLTQQER without any aromatic amirno acids or proline **(C)**.

## CONCLUSION

In this study we evaluated the fragmentation behavior of tryptic peptides in 213 nm UVPD. We see fragmentartion in time-scales feasible for large-scale bottom-up experiments, however fragmentation efficiency is comparrably low. Fragmentation seemed to be highly dependent on precursor properties, namely the charge density and the presence of aromatic amino acids (and proline). We see a possibility in the development of supplemental activation techniques using collision-based methods to help improve fragmentation. Also, use of an aromatic tag could increase photon absorption and possibly lead to increased spectral quality.

## ASSOCIATED CONTENT

### Supporting Information

The Supporting Information is available free of charge on the ACS Publications website.

Overview of PSMs found in Dual MS2 datasets and fragmentation analysis of the 4 PM dataset (PDF)

List of all analyzed *m/z* species (peptide sequence and charge state combination) (XLS)

## AUTHOR INFORMATION

### Notes

The Authors declare no competing financial interest.

## ACKNOWLEDGMENT

This work was supported by the Wellcome Trust (103139 and 108504) and the Deutsche Forschungsgemeinschaft (DFG, Gerrman Research Foundation, 329673113 and 426290502). The Wellcome Centre for Cell Biology is supported by core funding from the Wellcome Trust (203149).

---

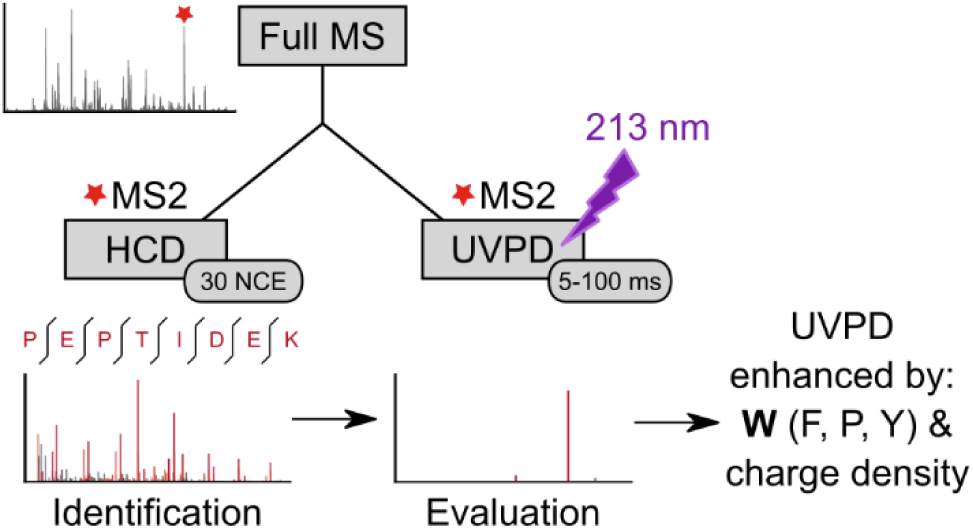

