## Supplemental Figures for "Ultraviolet Photodissociation of tryptic peptide backbones at 213 nm"

### Supporting Information: Ultraviolet Photodissociation of tryptic peptide backbones at 213 nm

#### List of supporting information

Table S1: List of all analyzed *m/z* species (peptide sequence and charge state combination).

Figure S1: Overview of unique peptide spectrum matches (PSMs) found in the Dual MS2 datasets

Figure S1: Fragmentation analysis of peptides from 4 PM dataset subjected to different UVPD excitation times and ETD, HCD and EThcD for comparison.

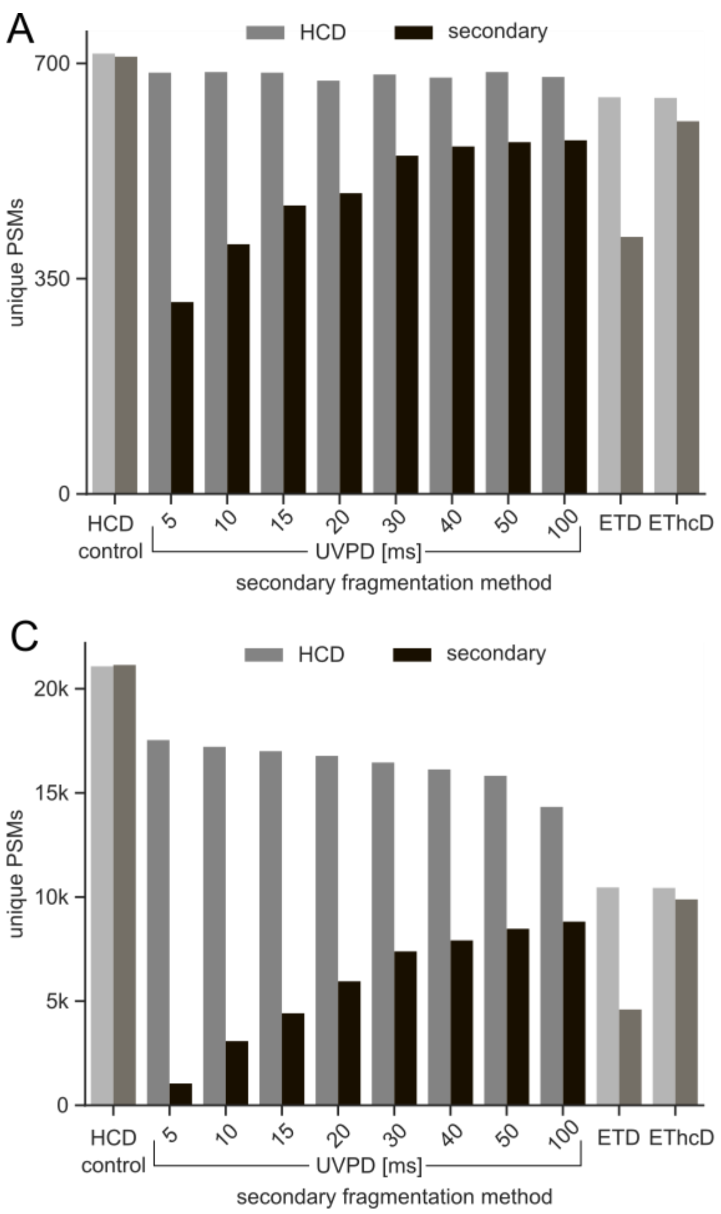


**Figure S1.** Overview of unique peptide spectrum matches (PSMs) found in the Dual MS2 datasets at 1% and 0.1% FDR for the 4 PM (**A**-**B**) and *E. coli* lysate (**C**-**D**) samples respectively. (**A, C**) Bar plots show the number of unique PSMs of the reference HCD scans in lighter color and the secondary fragmentation method scans HCD (positive control), UVPD 5-100 ms, ETD, and EThcD in darker color. (**B, D**) Venn diagrams showing the overlap of identifications from the HCD and UVPD scans from the HCD-UVPD Duals MS2 acquisitions.

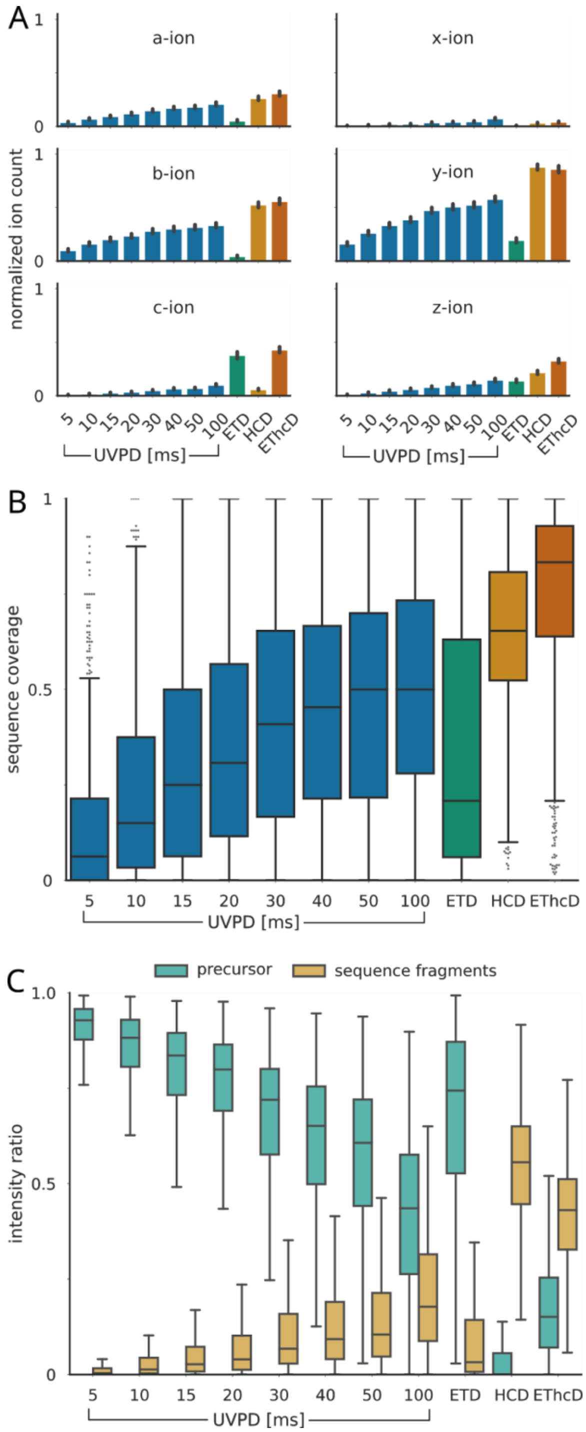


**Figure S2.** Fragmentation analysis of peptides subjected to different UVPD excitation times and ETD, HCD and EThcD for comparison. Data from 4 PM dataset. **(A)** Bar plots showing the sequence fragment ion type counts (normalized by peptide length). Error bars represent 0.95 confidence intervals. **(B)** Box plots showing sequence coverage of precursors. **(C)** Box plots of MS2 intensity ratios of remaining precursor and sequence fragments to total MS2 intensity. In all box plots whiskers extend to 1.5 interquartile range past the low and high quartiles.
